# Attraction of population receptive fields towards the attended locus is invariant to contrast

**DOI:** 10.1101/2025.07.01.662547

**Authors:** Sumiya Sheikh Abdirashid, Tomas Knapen, Serge O. Dumoulin

## Abstract

Attention enhances perception and spatial resolution. One mechanism by which attention achieves this is by attracting receptive fields towards the attended locus. The literature remains unclear on whether stimulus-driven parameters like contrast should alter this attraction. On one hand, Bayesian interpretation predicts changes in attraction due to contrast, whereas attention field models do not. Here we investigate whether stimulus contrast alters attention-driven attraction towards the attended locus. We used a demanding attentional task at fixation (0.1°) while mapping population receptive fields (pRFs) using either a low-contrast (5% Michelson contrast) or a high-contrast (80% Michelson contrast) checkerboard bar. Behavioral performance across conditions was matched. We show large and consistent differences in the amplitude of responses, but surprisingly, no difference in the amount of attraction towards the attended locus. The variance explained in this experiment was comparable to that of similar studies, which did observe pRF property changes, suggesting sufficient sensitivity to detect attraction towards the attended locus had it occurred at a similar magnitude. Our results cannot be explained by the Bayesian interpretation that predicts attention-based attraction effects varying with contrast. Instead, attention field models better account for our observations.

## Introduction

Visual attention improves perception by enhancing spatial resolution and contrast sensitivity (Moran and Desimone, 1985; Reynolds et al., 2000; Womelsdorf et al., 2006; Herrmann et al., 2010; Carrasco, 2011; Anton-Erxleben and Carrasco, 2013; Klein et al., 2014). For this reason, attention is often described as a zoom lens (Eriksen and St James, 1986). This attentional zoom lens is thought to be implemented on a neuronal level through gain changes (Moran and Desimone, 1985; Spitzer and Richmond, 1991; Brefczynski and DeYoe, 1999; Reynolds et al., 1999; Datta and DeYoe, 2009; Puckett and DeYoe, 2015; Fox et al., 2023), which result in the attraction of receptive fields towards the attended locus (Womelsdorf et al., 2006; Klein et al., 2014; van Es et al., 2018). This results in finer, denser sampling at the attended locus, improving spatial resolution (Reynolds and Heeger, 2009; Anton-Erxleben and Carrasco, 2013; Baruch and Yeshurun, 2014). It is well described how the position and precision of attention influence this attraction of receptive fields towards the attended locus (Klein et al., 2014; van Es et al., 2018; Liu et al., 2022; Tünçok et al., 2024; Abdirashid et al., 2025). What remains unclear is whether and how stimulus-driven properties, such as contrast, interact with attention-driven alterations in visual representation. Here, we investigate whether contrast alters the attraction of population receptive fields towards the attended locus.

Stimulus contrast alters neuronal firing rates (Albrecht and Hamilton, 1982; Sclar et al., 1990), the amplitude of responses (Boynton et al., 1999; Gardner et al., 2005; Marquardt et al., 2018), and some receptive field properties (Hunter and Born, 2011; Tsui and Pack, 2011; Butler et al., 2020). Additionally, contrast sensitivity is altered by attention, resulting in contrast response functions that are scaled additively and/or multiplicatively for attended stimuli compared to unattended stimuli (Luck et al., 1997; Martínez-Trujillo and Treue, 2002; Williford and Maunsell, 2006; Buracas and Boynton, 2007; Li et al., 2008; Herrmann et al., 2010; Foster and Ling, 2022). Stimulus contrast alters receptive field tuning, with lower contrast stimuli yielding larger receptive field sizes and lower neuronal signal-to-noise ratios (Kapadia et al., 1999; Sceniak et al., 1999, 2002; Kumano and Uka, 2012; Maloney and Clifford, 2015; Butler et al., 2020). In Bayesian terms, a lower contrast stimulus would create a broader likelihood distribution and result in an attention-driven, posterior response that is more heavily weighted towards the prior (i.e. the attention field). Thus, this Bayesian account would predict *more* attraction towards the attended locus for a lower contrast stimulus. In sum, both the Bayesian perspective and a broad body of literature suggest stimulus contrast should interact with receptive field attraction towards the attended locus.

However, there is an alternative body of literature which suggests otherwise. Many cortically tuned properties are contrast invariant, meaning their tuning functions are unchanged by stimulus contrast (Finn et al., 2007; Murray, 2008). In the case of orientation tuning, electrophysiology studies have found contrast invariant responses (Troyer et al., 2002; Finn et al., 2007), while in fMRI studies, the results are mixed (Maloney and Clifford, 2015; Liu et al., 2018; Butler et al., 2020). In some cases, as in macaque LOC, contrast invariant responses are observed for attended but not unattended stimuli (Murray and He, 2006). Attention field models, which describe the influence of attention on receptive field responses as a multiplication of Gaussians, predict that the amplitude of the stimulus-driven pRF should not alter attraction towards the attended locus (Womelsdorf et al., 2008; Reynolds and Heeger, 2009; Klein et al., 2014; van Es et al., 2018), though they do predict altered attraction if pRF size is changed or if a baseline is included in the attention field model. From both an attention field model perspective and based on the literature of contrast invariant cortically tuned visual features, an alternative hypothesis is that attraction of receptive fields towards the attended locus does not depend on contrast.

Thus, here we asked whether attention-driven population receptive field attraction is contrast dependent or contrast invariant. We used a demanding attentional task at fixation and a standard population receptive field mapping protocol with two different conditions: a low contrast stimulus (5% Michelson contrast) and a high contrast stimulus (80% Michelson contrast). As expected, there were reliable differences in BOLD amplitude between conditions, with the higher contrast condition having higher BOLD amplitudes. Because we collected more data for the low contrast condition, the variance explained was equated across conditions, as well as comparable to that of related studies, which did observe pRF property differences (Klein et al., 2014; Abdirashid et al., 2025). We found no significant contrast-dependent changes in pRF properties anywhere across the entire visual hierarchy. Our results suggest that attention-driven attraction of receptive fields towards the attended locus is contrast invariant and an attention field model interpretation, rather than a Bayesian framework, is best to describe these results.

## Methods

### Participants

Seven participants (ages 26 to 51, 4 female) were included in the final analysis for this experiment. Two were excluded from the final analysis due to excessive motion in the scanner and not completing the full session. All participants were screened before scanning to ensure MR compatibility. The study was conducted with informed consent from all participants. Ethical approval for this study was granted by the Human Ethics Committee of Vrije Universiteit Amsterdam (VWCE-2020-004R1). All participants had normal or corrected-to-normal visual acuity.

### Experimental Design

Visual stimuli were presented on an MR compatible Cambridge Research Systems LCD screen (69.84 × 39.29 cm, 1920 × 1080 pixels, 120-Hz refresh rate). In the scanner, the screen was located outside of the bore, and participants viewed it through a mirror mounted on the coil. The screen was 196 cm from the mirror. For training on the task outside of the scanner, we used a comparable setup with an identical screen, screen distance, and visual stimuli.

The visual stimuli were created using the PsychoPy Python package (Peirce, 2009) and the exptools2 wrapper for PsychoPy (https://github.com/VU-Cog-Sci/exptools2). A visual field mapping experimental design was used in combination with a demanding attentional task at fixation (Abdirashid et al., 2025).

Participants were instructed to fixate, ignore the checkerboard bar traversing the screen, and to conduct a task within the fixation circle. There were two mean-luminance fixation blocks at the start (20s) and end (30s) of every run. After the first 20s fixation block the attentional task began and the color dot stimulus was presented.

Within the 0.2° diameter fixation circle pink (RGB:1,0.35,0.87) and blue (RGB: 0.1,0.6,1) dots were presented on a mean luminance (22.1 cd/m^2^) background (Figure 1a). The dot positions and colors changed trial-by-trial, but remained within the fixation circle. Each trial consisted of a color dot stimulus presentation (375ms) and an interstimulus interval (375ms) (Figure 1b). On noise, or foil, trials the proportion of pink:blue dots was 50-50. Occasionally, there were target trials where the color dot stimulus became either more pink or more blue (i.e. proportion deviated from 50-50). The task was to detect these target trials. Roughly 1 in 12 stimulus presentations was a target, jitter was used to avoid predictability. This proportion detection task becomes more difficult the closer the target trials are to the 50-50 proportion. The target proportion was adjusted to ensure the difficulty and performance were matched across conditions and participants. All participants were trained on the task prior to scanning to avoid learning effects during scanning. A reaction time window of 1s was used to log correct responses.

**Figure 1:**
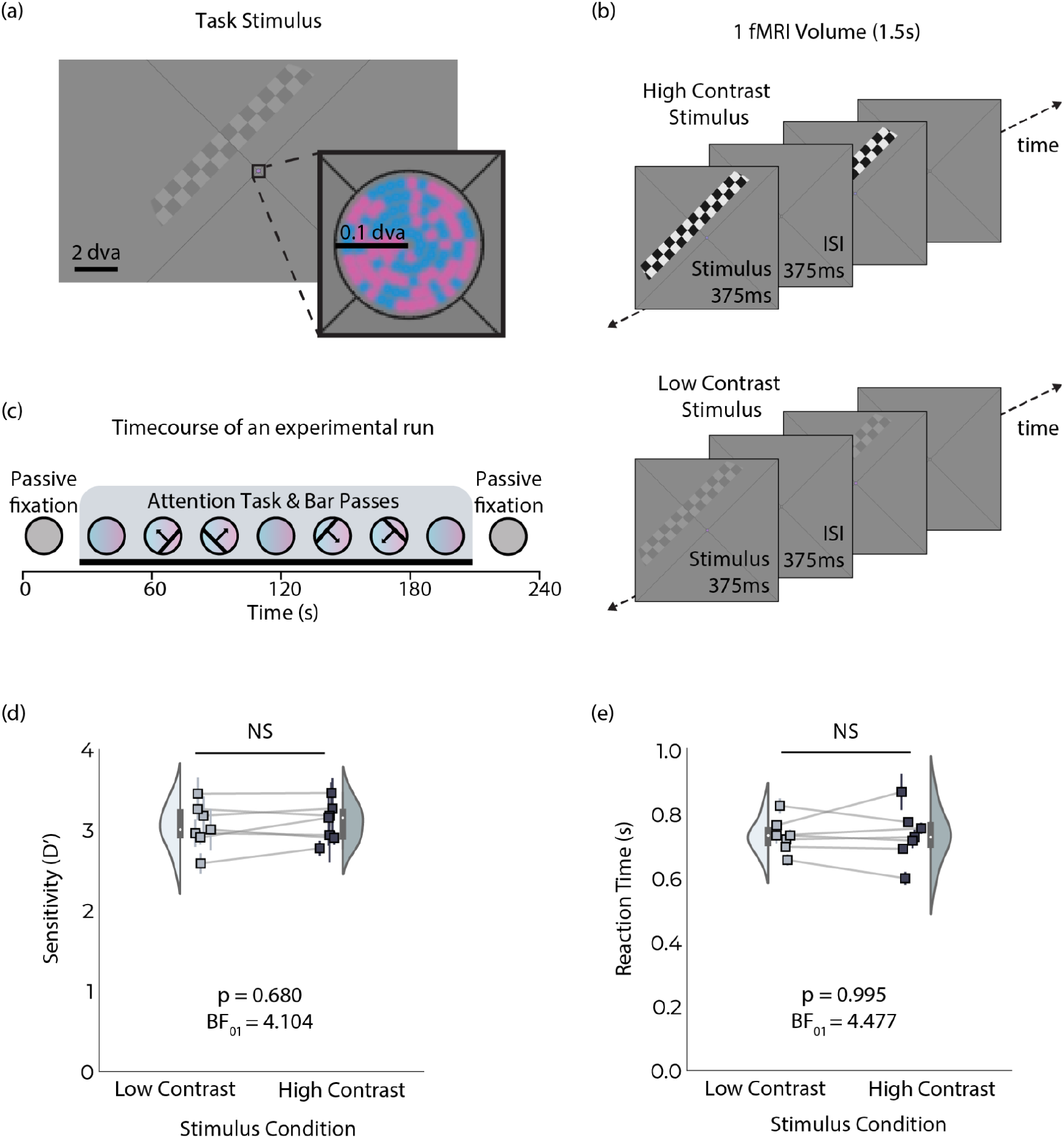
Experimental design and behavioural results. (a) An example stimulus from the low contrast condition. A color proportion discrimination task is performed within the fixation circle. A luminance contrast defined checkerboard bar traverses the screen. (b) Timing of stimulus presentation for the high contrast condition (80% Michelson contrast, top panel) and the low contrast condition (5% Michelson contrast, bottom panel). The checkerboard bar stimulus and the color dot task stimulus are presented for 375ms with a 375ms interstimulus interval. (c) Timecourse of an experimental run, the first 20s and the last 30s of each run only the mean luminance background and the fixation cross are presented. After 20s the task starts, and after another 10s the bar passes begin. (d) Sensitivity and reaction time (e) are indistinguishable across conditions.

15s after the start of the attentional task stimulus the visual field mapper started. A checkerboard bar stimulus (1.25° width) was used as a visual field mapper. It traversed the screen within a circular aperture with a diameter of ∼10° of visual angle. Two bar contrasts were used: low contrast (5% Michelson contrast, 20.3 cd/m^2^ black, 23.9 cd/m^2^ white) and high contrast (80% Michelson contrast, 4.02 cd/m^2^ black, 90.4 cd/m^2^ white). Bar contrasts were alternated on different runs. To compensate for the expected reduced signal-to-noise ratio in the low-contrast condition, we collected twice as many low-contrast runs. This ratio was derived from pilot experiments in which we estimated BOLD response contrast dependence. The bar swept the screen with two orientations (45°, 135°) and two motion directions (perpendicular to bar orientation), resulting in a total of four bar passes per run. The checkerboard pattern within the bar also moved parallel to the bar orientation. The bar motion was time-locked to the MRI acquisition and moved 0.625° each TR (1.5s). There was a 15s gap after every two bar sweeps, during which the attentional task continued. The bar sweeps were only presented within the duration of an attentional task. After the last bar sweep, there was an additional 30s of the task followed by 30s of mean luminance fixation.

### Independent visual field mapper

In addition to this experimental paradigm, standard visual field mapping was conducted for all participants in a separate session. In this session, subjects performed an easy (∼100% correct performance) color-change task at fixation, and the bar traversed the screen in 8 directions, including both cardinal and diagonal orientations. The visual stimuli and task further followed those outlined in the methods of Dumoulin and Wandell (2008) and Aqil et al., (2021).

### Statistics

In the two conditions of this experiment, participants were carrying out an identical task. To confirm that behavioural performance was comparable, we used both Bayesian and frequentist statistics. The discriminability index (D’) and reaction time were compared across subjects (Figure 1d,e). JASP Bayesian ANOVA was performed to test for evidence in favor of the null hypothesis. For this analysis, the dependent variable was the behavioural metric of interest (either D’ or reaction time), the task conditions were the fixed factors and subject and run were modeled as random effects variables (van Doorn et al., 2021). We also used a frequentist linear mixed model, with the same dependent variable, fixed effect variables, and random effect variables.

For statistical analysis of the fMRI data, first a Shapiro-Wilk test was used to test for normality. The data were not normally distributed so non-parametric statistical tests were selected for subsequent analyses. We performed a Wilcoxon signed-rank test to determine whether task pRF R^2^, amplitude, eccentricity and size (Figure 2-4) differed significantly between conditions. To correct for multiple comparisons the false-discovery rate (FDR) correction was used. To determine if there was any evidence in favour of the null hypothesis, we used Bayesian statistics. Since there is no multiple-testing correction with Bayesian statistics we opted for a Bayesian ANOVA with subject and ROI as random effects variables.

**Figure 2:**
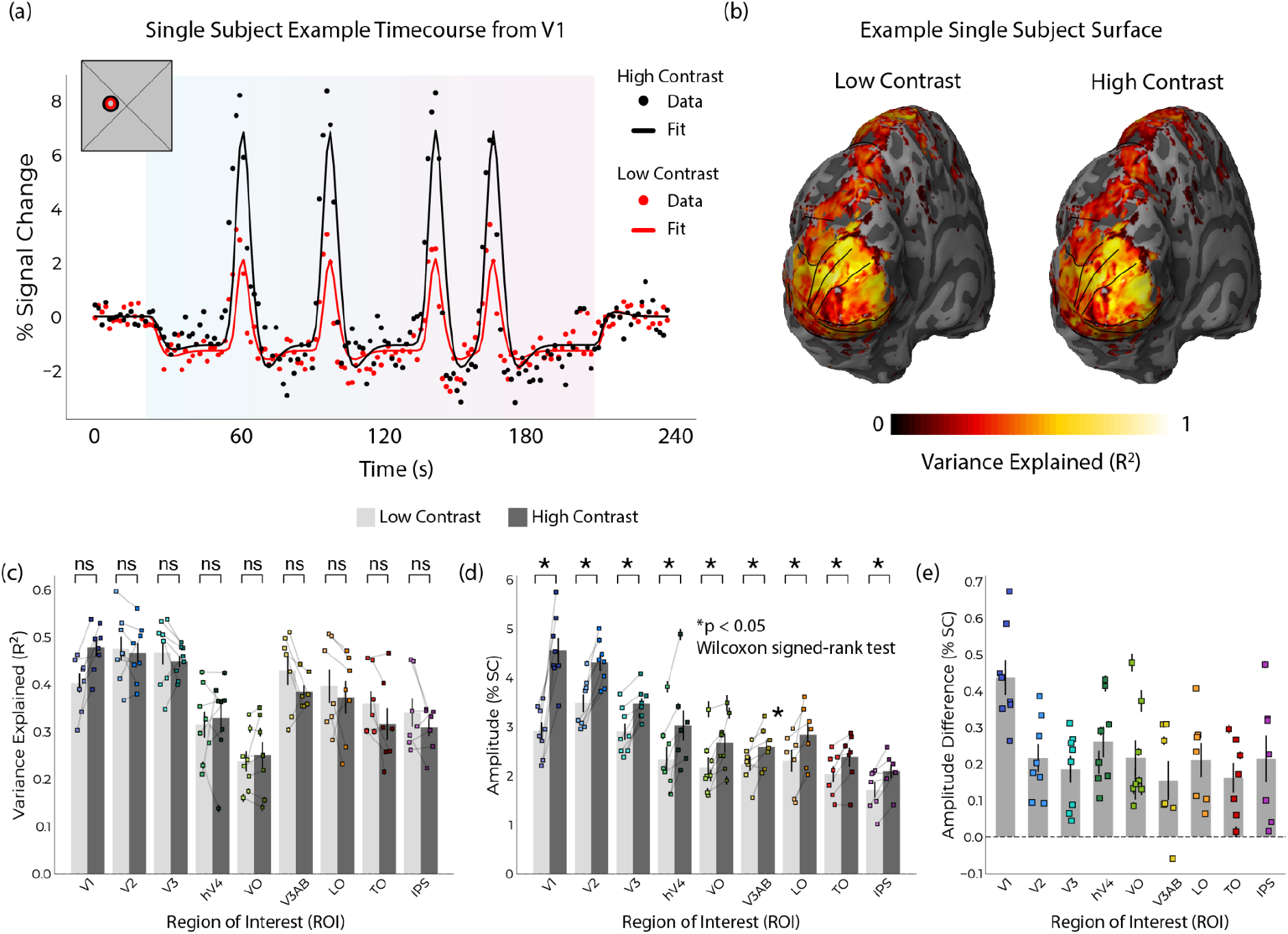
Variance explained is comparable across conditions, and our experimental design manipulates visual response amplitude. (a) Single subject example time course from a vertex in V1. The low contrast condition is shown in red and the high contrast condition is shown in black. Raw data is shown as points and the best-fitting model timecourse is shown as the solid line. The inset (upper left) is the pRF locations and sizes for each condition. (b) Variance explained of the pRF fits plotted on the two left hemisphere surfaces of the same example subject. On the left is the variance explained for the low contrast pRF fits and the right shows the variance explained for the high contrast fits. These maps are comparable. (c) Variance explained for each subject and each condition (shown as scatter) and group averages (shown as bar plots). (d) This design manipulates the amplitude of responses. The individual subject pRF amplitudes within each ROI are shown as individual points and group averages in the bar plots. The low contrast condition is shown in light gray and high contrast is shown in dark gray. (e) The difference of the amplitudes shown in (d) for each subject (scatter) and each visual area. Significant differences (FDR corrected p < 0.05) are indicated with an *. All error bars are SEM.

### Eye-tracking

All participants were trained before scanning on an equivalent set-up (same screen properties and distance). During this training and for most subjects during scanning gaze position was recorded using an Eyelink 1000 (SR Research, Osgoode, Ontario, Canada). The eye-tracker was calibrated using a 5-point procedure.

### MRI

All scans were acquired on a Phillips Achieva 7-T scanner with a 32-channel Nova Medical head coil. Foam padding was used to minimize head movement T1-weighted (T1w) MP2RAGE and T2-weighted (T2w) turbo-spin echo structural MRI scans were acquired at a resolution of 0.7mm isotropic (T1w: FOV = 220 × 220 × 200 mm^3^, matrix = 352 × 352 × 263, TR / TE = 6.2 ms / 3 ms, flip angle_1_ / flip angle_2_ = 5° / 7°, TR_MP2RAGE_/TI_1_/TI = 5500 ms / 800 ms/2700 ms, duration = 9 min 45 s; T2w: FOV = 245 × 245 × 184 mm^3^, matrix = 352 × 349 × 263, TR/TE = 3000/390 ms, TSE-factor = 182, duration = 7 min).

Functional MRI data was acquired with a 1.7 mm isotropic T2*-weighted gradient-echo 2D-EPI sequence containing 57 slices and 159 volumes (FOV = 216 × 216 mm^3^, matrix = 128 × 125, TR/TE = 1500ms / 22ms, flip angle = 53°). To avoid start-up magnetization transients, the first 12s (6 TRs) of acquisition were discarded. Every functional scan was followed by a top-up scan with the opposite phase-encoding direction for later susceptibility distortion correction. Each run had a scan duration of 330s (independent pRF mapper) or 239s (attentional task experiment).

### Preprocessing

Anatomical preprocessing followed a pipeline outlined by Heij and colleagues (Heij et al., 2023) (https://github.com/gjheij/linescanning). T1 and T2 images were combined using Freesurfer 7.2; this software was also used to reconstruct anatomical surfaces. Individual subject functional data was sampled to the individual subject surfaces using fMRIprep version 21 (Esteban et al., 2019). BOLD volumes were averaged to create one average timecourse, per vertex, per condition. These timecourses were linearly detrended and converted to % signal change using the blank fixation blocks at the beginning and end of each run.

### pRF & GLM fitting

Every experimental run had an event-related pRF design nested into a block design. The blocks were: 14 TRs (21s) passive fixation, 125 TR attention task & pRF mapping, and 20 TRs (31.5s) passive fixation (Figure 1c). pRF fitting was conducted on cut timecourses, excluding passive fixation blocks and any transient responses at the onset and offset of the task, as these could bias the baseline and amplitude estimates. Using the Python package prfpy (Aqil et al., 2021; Aqil and Knapen, 2023) (https://github.com/VU-Cog-Sci/prfpy), single Gaussian pRF fitting was carried out on the truncated timecourses (Dumoulin and Wandell, 2008).

To account for the blocks in the experimental design a GLM was used. This GLM was fit on the uncut timecourses. The two passive fixation blocks on either end of each run were used to estimate the BOLD baseline outside of the attentional task. The predictors in the design matrix were: the predicted timecourse from the pRF fitting, the attentional task (the TRs during which participants were doing the task), and two nuisance regressors at the timepoints of task onset and offset. All regressors were convolved with a canonical hemodynamic response function. For fitting GLM, least squares minimization was used.

To compare the amplitude of responses for the two conditions, the difference between the maximum and the minimum of the predicted timecourses was used.

### Visual field maps

Using the independent pRF session, visual field maps of eccentricity, polar angle, and pRF size were obtained for each participant. Using these maps, visual areas (V1, V2, V3, hV4, VO, LO, TO, V3AB, and IPS) were drawn following standards defined in the literature (Amano et al., 2009; Swisher et al., 2007; Wandell et al., 2005, 2007). Only vertices within the outlined regions of interest and surpassing an R^2^ threshold of 0.1 in all conditions were used in the final analysis.

## Results

### Behavioral performance is matched across conditions

In this experiment, we compared responses in two experimental conditions which were identical in the task being performed. Per design, behavioural performance for the two conditions was not significantly different (Figure 1d,e). The group averaged D’ and standard error was 3.050 ± 0.074 for the low contrast condition and 3.097 ± 0.087 for the high contrast condition. Reaction times were also comparable, with a group average of 0.733 ± 0.009 seconds for the low contrast condition and 0.729 ± 0.017 seconds for the high contrast condition. Individual subject behaviour can be found in supplementary materials (Supplementary Figures 1-3). Both frequentist and Bayesian statistical tests carried out indicate similar behavioural performance. A linear mixed model with subject and run as random effects factors showed no significant differences (p_D’_ = 0.680, p_RT_ = 0.995). A Bayesian ANOVA showed evidence in favour of the null hypothesis that the performance is similar (D’ BF_01_ = 4.104, reaction time BF_01_ = 4.477). Thus, we conclude that our experimental conditions were matched in their behaviour performance.

### The amplitude of responses significantly differs

To determine whether we successfully modulated the amplitude of responses with the mapper contrast, the minimum and maximum of the predicted timecourses were compared (Figure 2a,d,e). Differences in the amplitude of responses are shown in an example timecourse from a single vertex in V1 in Figure 2a. Amplitude differences were observed across subjects and along the visual hierarchy. As intended, this experimental design did significantly modulate the amplitude of responses (Figure 2d; Wilcoxon signed-rank test, FDR corrected *p*_*V1*_=0.03, *p*_*V2*_=0.03, *p*_*V3*_=0.03, *p*_*hV4*_=0.03, *p*_*V3A/B*_=0.84, *p*_*LO*_=0.06, *p*_*TO*_=0.04, *p*_*VO*_=0.03, *p*_*IPS*_=0.04). As expected from the increase in contrast invariance moving up the visual hierarchy (Buracas and Boynton, 2007; Roelofzen et al., 2025), the largest amplitude differences were seen in V1, with lower differences along the visual hierarchy.

### Variance explained for the conditions is matched

To account for the reduced signal-to-noise ratio in the low contrast stimulus, compared to the high contrast stimulus, we doubled the number of low contrast runs. To determine whether we correctly adjusted for this difference in signal-to-noise ratio, we looked at within-set variance explained (R^2^) from the pRF fitting (Figure 2b,c) on the data after averaging across runs. Variance explained did not significantly differ between the two conditions in any ROI (Figure 2c; Wilcoxon signed-rank test, FDR corrected *p*_*V1*_=0.14, *p*_*V2*_=0.84, *p*_*V3*_=0.86, *p*_*hV4*_=0.88, *p*_*V3A/B*_=0.84, *p*_*LO*_=0.94, *p*_*TO*_=0.84, *p*_*VO*_=0.84, *p*_*IPS*_=1.00).

Variance explained in this experiment is also the same or higher than experiments with similar designs combining an attention task and pRF mapping (Supplementary Figure 4b; Abdirashid et al., 2025; Klein et al., 2014; van Es et al., 2018).

### Stimulus contrast does not alter position preference

Attention is known to attract receptive fields towards the attended locus (Womelsdorf et al., 2006; Klein et al., 2014; van Es et al., 2018). Here, we investigated whether this attraction is modulated by stimulus contrast by directly comparing pRF position estimates from the low and high contrast conditions. To recapitulate, the Bayesian hypothesis predicts that decreasing stimulus contrast should increase the attraction of pRFs towards the fovea, resulting in lower eccentricity estimates across the visual system. The eccentricity maps for each condition plotted on the surface for one example participant (Figure 3a) are comparable. The pRF position estimates did not significantly differ across participants and along the visual hierarchy (Figure 3b, c). There were no significant differences between the two conditions (Wilcoxon signed-rank test, FDR corrected *p*_*V1*_ = 0.88, *p*_*V2*_ = 0.56, *p*_*V3*_ = 0.56, *p*_*hV4*_ = 0.14, *p*_*VO*_ = 0.56, *p*_*V3A/B*_ = 0.56, *p*_*LO*_ = 0.94, *p*_*TO*_ = 1.00, *p*_*IPS*_ = 0.88). We also conducted Bayesian statistics to probe if there is evidence in favour of the null hypothesis that there is no significant difference between these two contrast conditions. Bayes Factor (BF_01_) values between 3 and 10 indicate moderate evidence in favour of the null hypothesis (van Doorn et al., 2021). Indeed, in a Bayesian ANOVA with subject and ROI as random effects variables, we obtained a BF_01_ = 5.084, indicating moderate evidence in favour of the null hypothesis.

**Figure 3:**
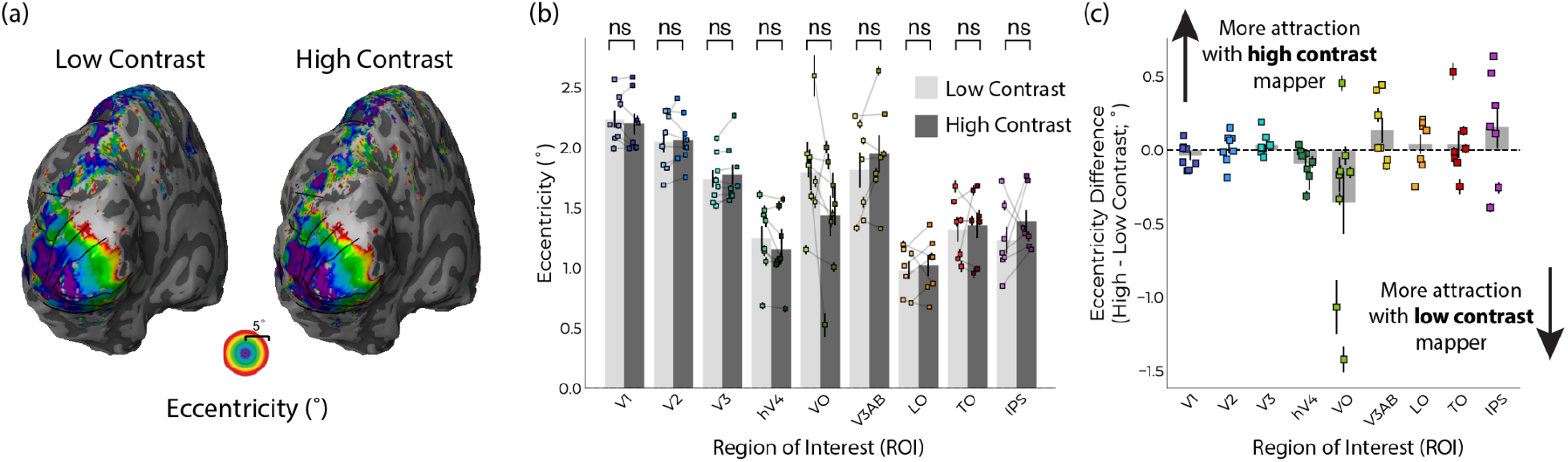
Stimulus contrast does not modulate pRF position. (a) pRF eccentricity plotted on two left hemisphere surfaces for one example subject. The left shows fitted eccentricity values for the low contrast condition, while the high contrast eccentricity values are shown on the right surface. Black lines indicate visual area boundaries. (b) and (c) eccentricity does not significantly (NS) differ across conditions. Bar heights in (b) and (c) show the mean across participants, per visual region; data points show the R2-weighted mean for each participant’s visual region; error bars are SEM, corrected for volume-to-surface upsampling.

There were also no statistically significant differences in pRF size across individuals and along the visual hierarchy (Supplementary Figure 5). There was also moderate evidence in favour of the null hypothesis (Bayesian ANOVA with subject and ROI as random effects variables BF_01_ = 4.931).

## Discussion

In this study, we sought to investigate whether the attraction of receptive fields towards the attended locus depends on the contrast of stimulus-driven responses. We compared two conditions identical in attentional task (Figure 1a), behavioural performance (Figure 1d,e), and model fit quality (Figure 2b,c). These two conditions differed only in the contrast of the mapping stimulus: either low contrast (5% Michelson contrast) or high contrast (80% Michelson contrast). As intended, we found significant modulation of the amplitude of responses across the visual hierarchy. Using a linear Gaussian pRF model, we did not see any significant differences in pRF size or pRF position. Using Bayesian analyses, we further found moderate evidence in favour of the null hypothesis that there are no differences in pRF size or position. The variance explained in this experiment (Figure 2c) was comparable to that of similar experiments (Klein et al., 2014; van Es et al., 2018), which did observe pRF position changes. Therefore, if there were differences in pRF parameters in the same range of magnitude as these experiments (∼0.5°), we would have had the signal to observe them. In this study, the attraction of pRFs towards the attended locus was invariant to contrast.

Here, we investigated altering the amplitude of an unattended, task-irrelevant stimulus. We chose a demanding behavioural task with a difficulty that is titratable instead of a fixation dot color task commonly used in pRF mapping experiments. This allowed us to limit exogenous attention differences resulting from the different bar contrasts. Indeed, behaviour was matched between the two conditions. This design allowed us to isolate any interactions between a consistent attention field and a changing stimulus-driven amplitude. Perhaps for stimulus-driven changes to alter the attraction of receptive fields, they need to be task-relevant or spatially overlapped with the attention field. Recent work from our lab does demonstrate that the attention field does not operate uniformly, and that it interacts with pRFs it has an overlap with (Abdirashid et al., 2025).

It is possible that contrast-dependent changes in receptive field attraction towards the attended locus could occur on a scale that we did not have the resolution to detect. Other designs which observed pRF property changes in the range of ∼0.5° manipulated the position, attention and/or size of attention (Klein et al., 2014; van Es et al., 2018; Abdirashid et al., 2025). This indicates that even if there was stimulus-driven interaction with attentional attraction, this does not occur in the same range of magnitude as manipulating voluntary attention. Furthermore, stimulus-driven contrast has the largest effect in V1; however, attention effects are largest further up the visual hierarchy. This potentially limits the population of neurons in which this interaction could be visible. It is possible these interactions could be visible in a smaller sub-population of neurons. For example, in neurophysiology experiments, orientation tuning is contrast invariant (Troyer et al., 2002; Alitto and Usrey, 2004; Finn et al., 2007) while in BOLD fMRI contrast alters orientation tuning bandwidth (Maloney and Clifford, 2015; Butler et al., 2020).

From a model perspective, a Bayesian model predicts that altering stimulus-driven contrast (i.e. likelihood) with a consistent attention field (i.e. the prior) should alter the posterior (i.e. the attention-driven pRF). However, a Gaussian attention-field model predicts that the amplitude of the stimulus-driven receptive field should not alter the attraction of population receptive fields towards the attended locus. Here we altered the gain of responses, but observed no differences in attention-driven pRF properties. Our observations are more in keeping with an attention-field model perspective.

## Supplementary Materials

**Supplementary Movie 1:** High-contrast experimental stimulus movie.

**Supplementary Movie 2:** Low-contrast experimental stimulus movie.

**Supplementary Figure 1:**
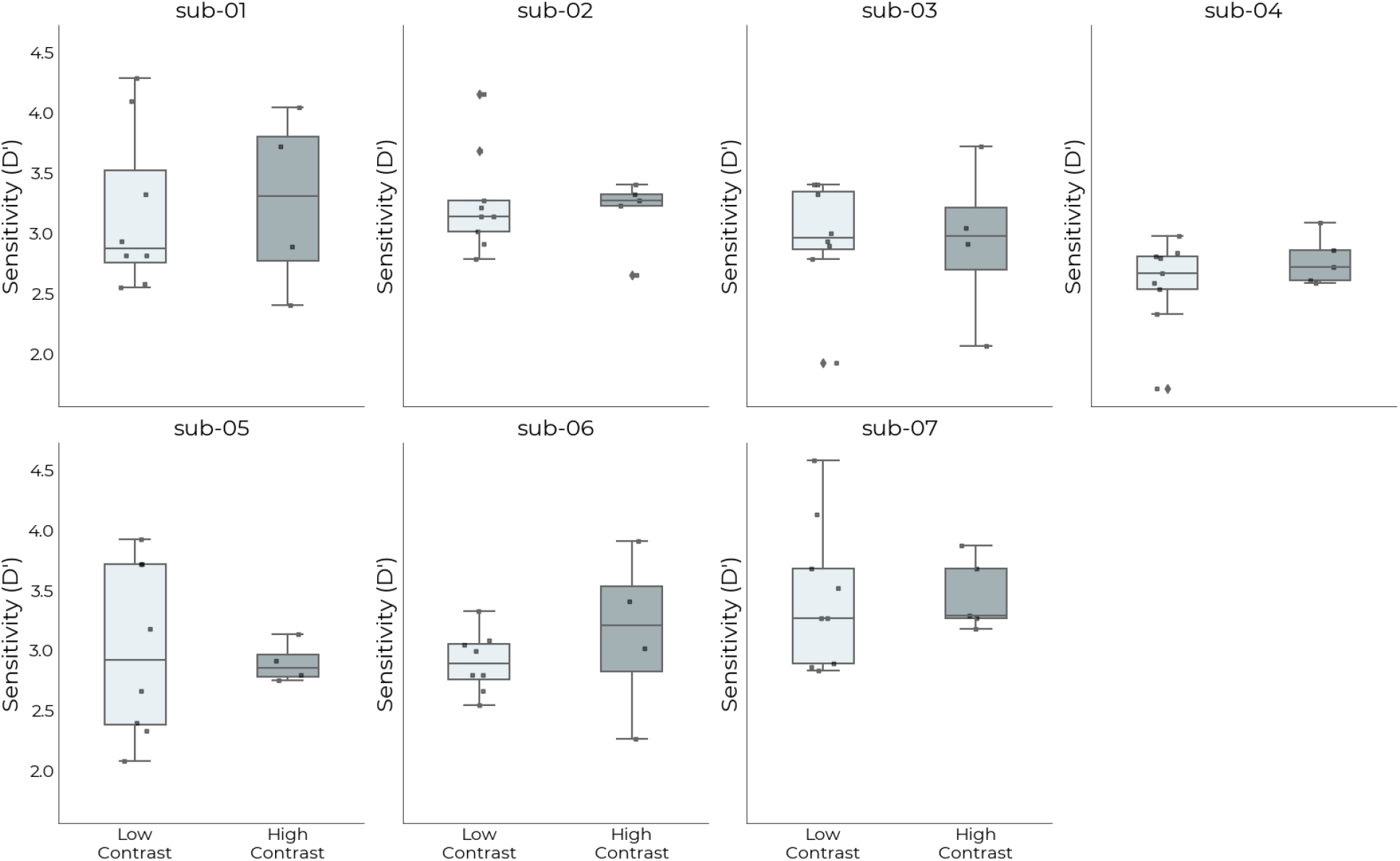
Individual subject sensitivity (D’). Paired boxplots indicating sensitivity to the demanding attentional task at fixation. Light gray color indicates low contrast condition, while dark gray denotes high contrast condition. Individual points show D’ per run. The low contrast condition had more runs than the high contrast condition.

**Supplementary Figure 2:**
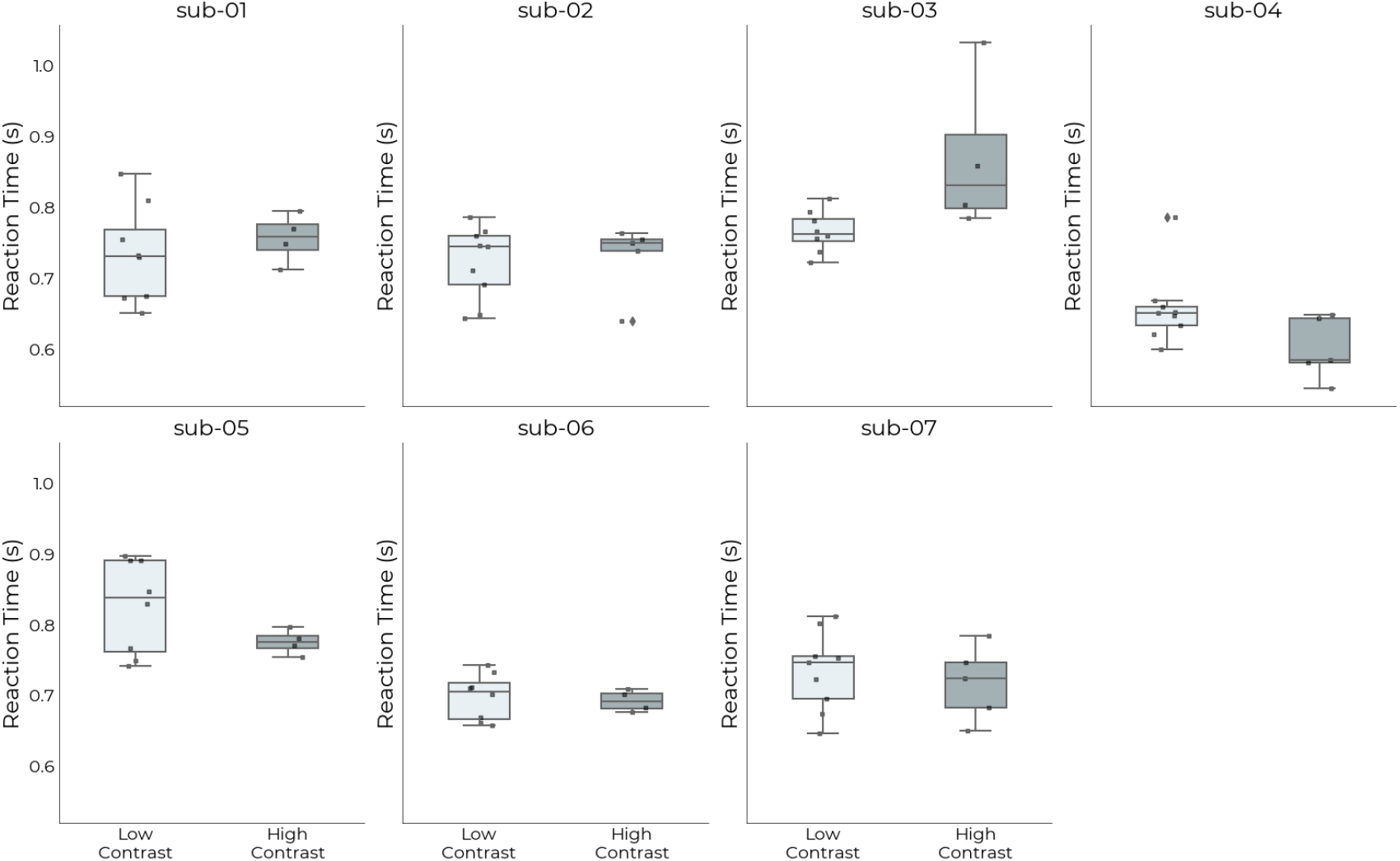
Individual subject reaction time. Paired boxplots showing reaction time in seconds to the attentional task at fixation. Participants had a maximum response window of 1s for the response to be counted. Light gray color indicates low contrast condition, while dark gray denotes high contrast condition. Individual points show reaction time per run.

**Supplementary Figure 4:**
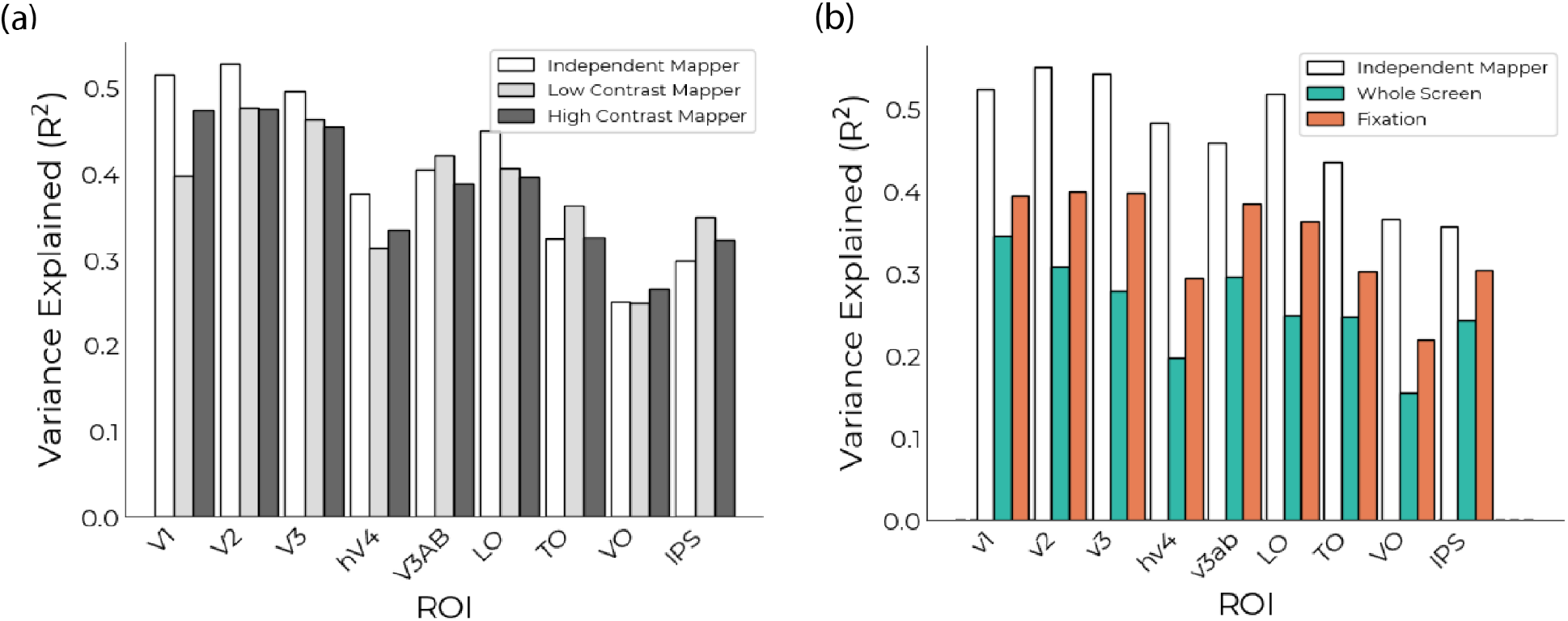
Variance explained (R^2^) in this experiment is comparable or higher than previous experiments which are able to observe eccentricity differences across conditions. (a) Variance explained from this experiment plotted with variance explained of the independent mapper (white). Low contrast variance explained for each visual area is shown in light gray and high contrast variance explained is shown in dark gray. (b) Variance explained from (Abdirashid et al., 2025), where they observed a significant eccentricity difference between experimental conditions.

**Supplementary Figure 5:**
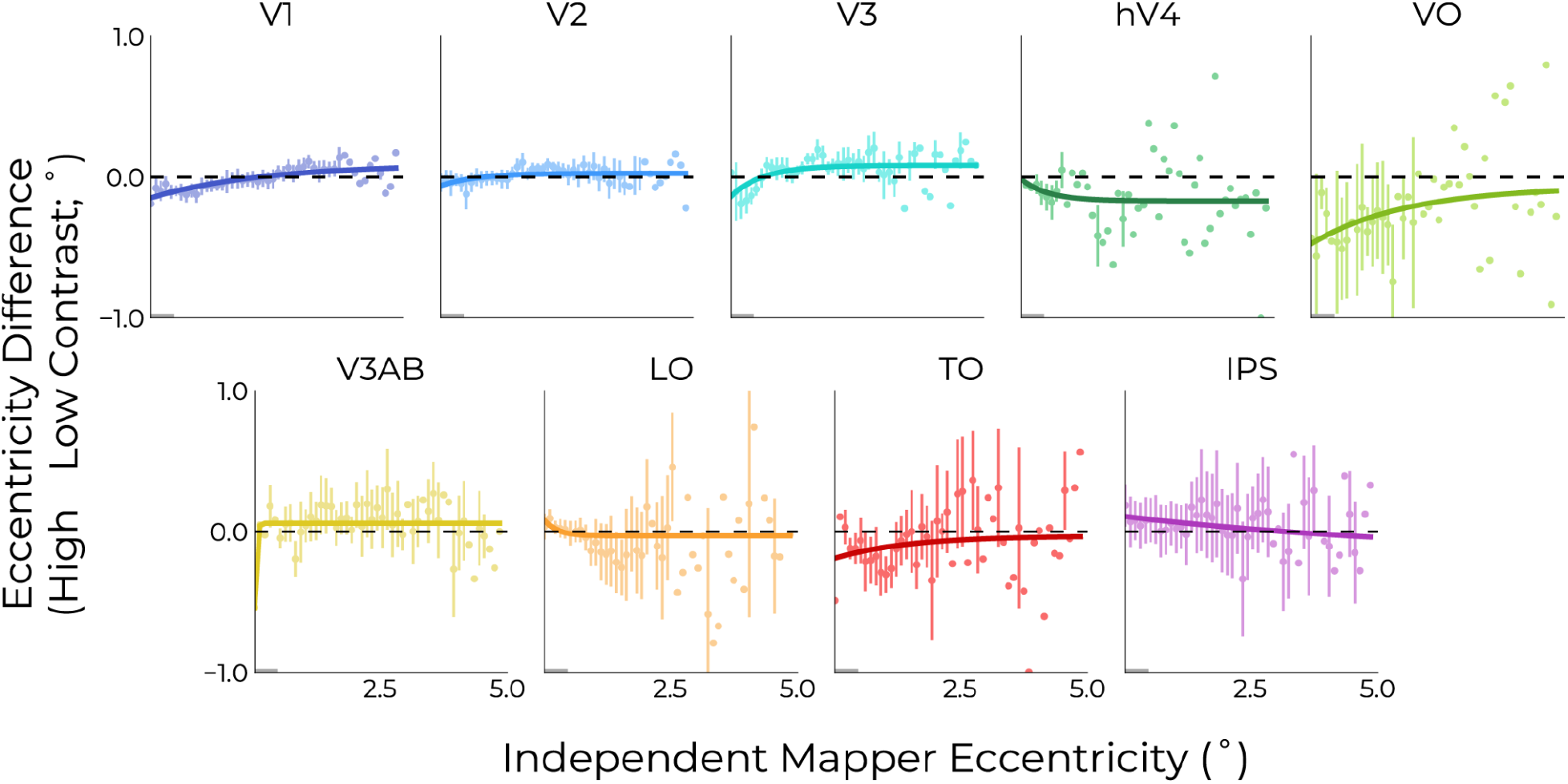
Eccentricity difference along binned eccentricity from the independent mapper. These values are first averaged within each bin for each participant and then averaged as a group. Error bars are SEM.

**Supplementary Figure 6:**
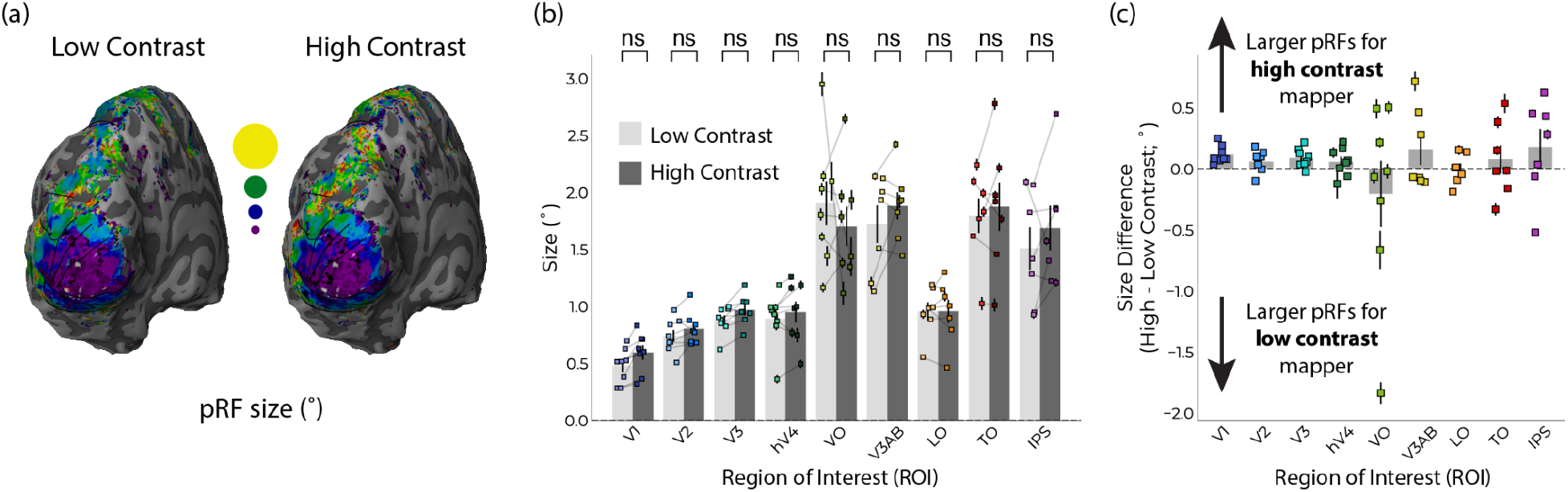
Stimulus contrast does not modulate single-Gaussian pRF size. (a) pRF size plotted on two left hemisphere surfaces for one example subject. The left shows fitted pRF size values for the low contrast condition, while the high contrast pRF size values are shown on the right surface. Black lines indicate visual area boundaries. (b) and (c) pRF size does not significantly (NS) differ across conditions. Bar heights in (b) and (c) show the mean across participants, per visual region; data points show the R2-weighted mean for each participant’s visual region; error bars are SEM, corrected for volume-to-surface upsampling.

